# High-content quantitative high-throughput screening identifies a cell cycle-associated signaling cascade that regulates a multienzyme metabolic assembly for glucose metabolism

**DOI:** 10.1101/2022.06.10.495654

**Authors:** Danielle L. Schmitt, Patricia Dranchak, Prakash Parajuli, Dvir Blivis, Ty Voss, Casey L. Kohnhorst, Minjoung Kyoung, James Inglese, Songon An

## Abstract

We have previously demonstrated that human liver-type phosphofructokinase 1 (PFK1) recruits other rate-determining enzymes in glucose metabolism to organize multienzyme metabolic assemblies, the glucosomes, in human cells. However, it has remained largely elusive how glucosomes are reversibly assembled and disassembled to functionally regulate glucose metabolism in human cells. We developed a high-content quantitative high-throughput screening (qHTS) assay to evaluate the impact of small molecule libraries on the formation of PFK1-mediated glucosome assemblies from stably transfected HeLa Tet-On cells. Initial qHTS with a library of pharmacologically active compounds directed following efforts to kinase-inhibitor enriched collections. Consequently, three active compounds that were known to inhibit cyclin-dependent kinase 2, ribosomal protein S6 kinase and Aurora kinase A, respectively, were identified and further validated under high-resolution fluorescence single-cell microscopy. Subsequent knockdown studies using small-hairpin RNAs confirmed an active role of Aurora kinase A on the formation of PFK1 assemblies in HeLa cells. Importantly, all the identified protein kinases here have been investigated as key signaling nodes of one specific cascade that controls cell cycle progression in human cells. Collectively, our qHTS approaches unravel a cell cycle-associated signaling cascade that regulates the formation of PFK1-mediated glucosome assembly in human cells.

## Introduction

Enzymes in glycolysis have been identified to form a multienzyme metabolic assembly at subcellular levels in a variety of species (1,2). Particularly, glycolytic enzymes in human erythrocytes are shown to colocalize with a membrane-bound band 3 protein on the inner cell membrane (3). In addition, human liver-type phosphofructokinase 1 (PFK1) is demonstrated to form a multienzyme metabolic assembly, namely the glucosome, in the cytoplasm of several human cell lines by recruiting other enzymes that catalyze rate-determining steps in glycolysis as well as in gluconeogenesis (4). Recently, phase separation-mediated mechanisms for metabolic assemblies have been revealed to explain the formation of glucosome assemblies in human cells (5) as well as glycolytic bodies in yeast cells (6). Nonetheless, it has yet to be elucidated, particularly in human cells, which regulatory mechanisms control glucosome assemblies and their dynamics for their functional contributions to cellular metabolism and beyond.

Meanwhile, glucose metabolism has been shown to functionally associate with the cell cycle in yeast and mammalian cells (7,8). Enzyme expression and activity of phosphofructokinase-2/fructose-2,6-bisphosphatase (PFKFB3) are significantly increased during the G1 phase, which in turn activates PFK1 to upregulate glycolysis (9,10). Similarly, an increased expression of hexokinase 2 leads to the promotion of glycolysis during the G1/S transition (11). At the same time, interactions between cyclin dependent kinases (CDKs) and their associated cyclins are essential to regulate subcellular locations, expression levels, and/or enzymatic activities of glycolytic enzymes, including PFK1 and pyruvate kinase muscle isoform 2, for cell cycle progression (10,12-14). Importantly, we have recently revealed that multienzyme glucosome assemblies, that regulates glucose flux at subcellular levels (4,5,15), dynamically oscillate in space and time during a cell cycle (16). However, it has remained elusive which regulatory mechanisms ensure tight control over glucosome assemblies and their dynamics during cell cycle progression.

In addition, microscopy-based phenotypic screening assays have been advantageous for investigating regulatory mechanisms of various cellular processes in human cells (17-19). Traditional high-throughput screening methods using small molecule libraries at single concentrations are not designed to capture a full pharmacological response profile, therefore potentially missing biologically relevant phenotypes accompanied by biphasic or bell-shaped concentration responses. Accordingly, quantitative high-throughput screening (qHTS) methods have been developed to pharmacologically evaluate small molecule libraries in a multi-point titration, producing concentration-response curves accompanying each small molecule (20,21). In fact, the qHTS assays have been successfully employed particularly when cell-based assays report protein translocation to or from cellular compartments (22) or changes of protein expression levels (23-25). Therefore, we sought to systematically dissect cellular mechanisms that regulate the reversible formation of a multienzyme glucosome assembly by employing qHTS technology.

In this work, we first developed a cell-based phenotypic qHTS assay using stably transfected HeLa cells that express PFK1 with a monomeric form of enhanced green fluorescent protein (PFK1-mEGFP) as a glucosome marker. After screening more than 4200 molecules, we identified three pharmacological molecules that are known to inhibit CDK2, ribosomal protein S6 kinase (RSK) and Aurora kinase A (AURKA), respectively. Our subsequent knockdown studies using small-hairpin RNAs (shRNAs) confirmed the importance of AURKA in regulation of PFK1-mediated glucosome formation in HeLa cells. Importantly, all three protein kinases we identified here have been characterized to form one specific signaling cascade (i.e., RSK-CDK2-AURKA) that plays an important role during cell cycle progression in human cells (26-29). Therefore, we demonstrate that PFK1-mediated glucosome assemblies are spatiotemporally regulated by the RSK-CDK2-AURKA signaling cascade during a cell cycle of human cells.

## Materials and Methods

### Materials

A plasmid expressing PFK1 conjugated with a monomeric enhanced green fluorescent protein (mEGFP) was prepared previously (4). Note that mEGFP contains three point mutations, A206K, L221K, and F223R, which prevent oligomerization of EGFP in cells (30,31). Small molecules used in this study were dissolved in DMSO (Sigma) and used at the concentrations indicated, that include resveratrol (Sigma), clemastine fumarate (Sigma), ARP101 (Tocris), 1,10-phenanthroline (Acros Organics), triflupromazine hydrochloride (Sigma), kenpaullone (Sigma), calmidazolium chloride (Sigma), PAC-1 (Sigma), SU9516 (Tocris), SL-0101-1 (Tocris), palbociclib (Sigma), BS-181 (Tocris), PF-03814735 (Sigma or Cayman), hesperadin (Tocris), bisindolylmaleimide II (BIM II, Cayman), and BIM X (Cayman).

### Generation of stably transfected HeLa-T-PFK1-mEGFP cell line (HeLa-T-PFK1G)

A HeLa-T-PFK1-mEGFP stable cell line (hereafter, HeLa-T-PFK1G) was generated using the HeLa Tet-On® 3G inducible expression system (Clontech) according to the manufacturer’s protocol. Briefly, the cDNA of PFK1-mEGFP was amplified by PCR and cloned into pTRE3G vector (Clontech). The pTRE3G vector containing PFK1-mEGFP was then co-transfected into HeLa Tet-On 3G cells (Clontech) along with a linear selection marker of hygromycin using Xfect (Clontech). These cells were cultured in a medium containing 200 µg/mL G418 sulfate (Santa Cruz Biotechnology). After stabilizing and splitting confluent cells, clonal selection was begun by introducing 100 µg/mL hygromycin B (Invitrogen). Cells were monitored daily, and the medium was exchanged until the formation of drug-resistant colonies was observed. Isogenic harvesting of colonies was achieved using sterilized cloning disks saturated with a Trypsin:EDTA solution (Corning, Cat# 25-053-Cl) to gently remove and seed them in a fresh medium. Colonies were then screened for PFK1-mEGFP expression in the absence and presence of 1 µg/mL doxycycline under fluorescence microscopy. A colony with the highest expression level (i.e., brightest mEGFP fluorescence) was selected. We subsequently validated that our stable cell line has a reproducible expression efficiency of ∼85-90% of cell population in response to doxycycline. HeLa-T-PFK1G cells were maintained in Roswell Park Memorial Institute 1640 (RPMI 1640, Mediatech, Cat# 10-040-CV) or Dulbecco’s Modified Eagle Medium (DMEM with high glucose, Gibco, Cat# 11965-092) supplemented with 10% dialyzed fetal bovine serum (FBS, Atlanta Biologicals, 12-14 MWCO, Cat #S12850) under the antibiotic selection of 200 µg/mL G418 sulfate (Santa Cruz Biotechnology) and 100 µg/mL hygromycin B (Invitrogen), along with either 50 µg/mL gentamycin sulfate (Corning) or 1% penicillin-streptomycin (Gibco) for maintenance.

### Transient transfection

Parental HeLa (ATCC CCL-2) cells were acquired from the American Tissue Culture Collection (ATCC, Manassas, VA) and maintained in RPMI 1640 supplemented with 10% dialyzed FBS and 50 µg/mL gentamycin sulfate in a HeraCell CO_2_ incubator (37 °C, 5% CO_2_, and 95% humidity). Briefly, HeLa cells were plated on either glass-bottomed 35 mm Petri dishes (MatTek) or 8-well chambers (LabTek) in RPMI 1640 supplemented with 10% dialyzed FBS without antibiotics. The following day, cells were transfected with PFK1-mEGFP using Lipofectamine 2000 (Invitrogen, Cat# 11668019) in Opti-MEM I (ThermoFisher, Cat# 11058-021) and allowed to incubate in a HeraCell CO_2_ incubator (37 °C, 5% CO_2_, and 95% humidity). Approximately 5 hours later, the medium was exchanged for a fresh growth medium and incubated overnight in the CO_2_ incubator.

### High-resolution fluorescence live-cell imaging

A day before imaging, HeLa-T-PFK1G cells in RPMI 1640 supplemented with 10% dialyzed FBS, 200 µg/ml G418 sulfate, and 100 µg/ml hygromycin B, were plated in the presence of 1 µg/mL doxycycline to induce the expression of PFK1-mEGFP. On the day of imaging, cells were washed three times for 10 minutes with an imaging solution (20 mM HEPES (pH 7.4), 135 mM NaCl, 5 mM KCl, 1 mM MgCl_2_, 1.8 mM CaCl_2_ and 5.6 mM glucose) and allowed to adjust to ambient temperature for ∼45-60 minutes prior to small molecule treatments. Alternatively, parental HeLa cells were treated with small molecules and incubated at 37 ^o^C for a given time after ∼24 hours following transient transfection of PFK1-mEGFP or PFK1-mCherry.

All samples were then imaged at ambient temperature (∼25 ^o^C) with a Nikon 60x objective lens (NA = 1.45, CFI Plan Apo TIRF) using a Photometrics CoolSnap EZ monochrome CCD camera at 1×1 binning on a Nikon Eclipse Ti-inverted C2 confocal microscope. Epifluorescence imaging was carried out using a set of Z488/10-FC cleanup, HC TIRF Dichroic and 525/50-HC emission filter from Chroma Technology. Images were acquired using NIS Elements (Nikon) and images were analyzed using an ImageJ freeware (National Institutes of Health).

### Compound libraries

We performed qHTS assays with two sets of chemical libraries that the National Center for Advancing Translational Sciences (NCATS) acquired: i) the library of pharmacologically active compounds (LOPAC, 1280 compounds, Sigma-Aldrich), and ii) the kinase inhibitor-enriched libraries (KIEL, 2958 compounds) that are composed of the Protein Kinase Inhibitor set (32,33), the Roche Kinase Inhibitor Library (34), and NCATS mechanism interrogation plates (35). Library preparation and management were performed as previously described (36). Briefly, compounds were plated into 384-well plates (Greiner Bio-One) comprising a 11-point intra-plate titration with a serial dilution of 1:3 in DMSO ranging from 10 mM to 169 nM. Plates were reformatted in quadrants into 1536-well format using an Evolution P3 system (PerkinElmer).

### Quantitative high-content high-throughput screening (qHTS) assay

HeLa-T-PFK1G cells were plated in 1536-well black, clear bottom, low base microwell plates (Aurora Microplates, Inc.) at a density of 400 or 600 cells/well in DMEM supplemented with 10% dialyzed FBS, 1% Pen-Strep, and 1 µg/mL doxycycline. Cells were incubated at 37 °C, 5% CO_2_, and 95% humidity for 16 hours. Please note that a qHTS assay was scaled to 4 µL volumes in 1536-well format in a semi-automated manner.

An initial screen of the LOPAC library (Sigma, 6.6 mM – 10 mM, 7-point 1:3 inter-plate titration) was performed. 23 nL of compound were transferred to respective wells with a pintool (Kalypsys). The plates were incubated overnight at 37 °C, 5% CO_2_, and 95% humidity. The following day, resveratrol was dispensed at a final concentration of 200-250 µM using a Mosquito Dispenser (TTP Labtech) as a positive control on each assay plate. The plates were then incubated at room temperature for 4 hours. Following incubation, cells were fixed with 3.5% paraformaldehyde and nuclei were stained with a 1:1000 dilution of Hoechst dye (Invitrogen) for ∼20 minutes. Cells were washed twice with 1x PBS with an EL406 microplate washer-dispenser (BioTek Instruments) and imaged using an INCell Analyzer 2000 (GE Healthcare Life Sciences) that is equipped with a Nikon 20x objective lens (NA = 0.45, Plan Fluor, Elwb Corr Collar = 0-2.0, CFI/60). Imaging of PFK1-mEGFP was done using a FITC channel (490/20x excitation, 525/36m emission) while imaging of the Hoechst dye was performed using a DAPI channel (350/50x excitation, 455/50m emission). Laser autofocus with offsets of 10.0 and 12.0 with 0.100 and 0.500 exposures, respectively, and 2×2 camera binning were used in a horizontal serpentine acquisition motion to collect one field of images from each well.

A target class-directed second qHTS using the KIEL libraries enriched for kinase inhibitors included the Protein Kinase Inhibitor set (32,33), the Roche Kinase Inhibitor Library (34), and NCATS mechanism interrogation plates (35). These libraries were tested at 10 mM, 11-point 1:3 intra-plate titrations as above and compounds were transferred in a final concentration range of 974 pM to 57.5 µM. Control compounds of SU9516, kenpaullone, and olomoucine, based on the LOPAC screening result, were used in an 8-point 1:3 titration (57.5 µM to 26.3 nM final assay concentrations) as intraplate controls. Approximately 24 hours later cells were fixed and imaged with an INCell Analyzer 2000 or 2200 (GE Healthcare Life Sciences) as described above.

Lastly, selected compounds, SU9516, SL-0101-1, PF-03814735 and kenpaullone, were further evaluated at 10 mM, 16-point 1:2 intra-plate titrations as above in a final concentration range of 1.8 nM to 57.5 µM. BIM II and BIM X, which are known for producing artifactual fluorescence signals (24), were also tested in a final concentration range of 1.8 nM to 57.5 µM. Approximately 24 hours later cells were fixed and imaged with an ImageXpress (Molecular Devices) using a Nikon 40x objective lens (NA = 0.95, Plan Apo Lambda). Four images per well were collected at 1×1 camera binning. Image analysis was performed using Columbus software (PerkinElmer).

### Data analysis of qHTS

Cell images were processed in the GE Workstation software and analyzed with a multitarget analysis of fluorescence. Top-hat segmentation of a DAPI channel was used to identify nuclei with a minimum area 100 µm^2^ and sensitivity of 40 settings. Multiscale top-hat segmentation of a FITC channel was used to identify cells with a characteristic area of 200 µm^2^ and sensitivity of 70 settings. This allowed for identification of fluorescent cells from background or dead cells. Following cell segmentation, the same FITC channel was analyzed with multiscale top-hat segmentation to identify PFK1-mEGFP assemblies with a size range of 0.2 – 2 µm^2^, number scales of 1, process parameters of 1 pixel, sensitivity of 40, and inclusions in the cytoplasm. A filter of object area, that is, the area of PFK1-mEGFP assemblies, below 30 µm^2^ and an advanced sensitivity range of 1.5 were also applied. A decision tree thresholding was developed to identify cells showing PFK1 assemblies with three nodes. The start node required a cell intensity above 110 GL, the second node required a PFK1 assembly count above 1, and the third node required a total area of PFK1 assemblies above 5 µm^2^. Ultimately, this allowed for identification of assembly-positive cells over assembly-negative cells. The total number of cells with a total area of PFK1 assemblies above 5 µm^2^ (most stringent parameter) and the total number of cells with any PFK1 assemblies were calculated with the above algorithm and normalized by plate to the 57.5 µM kenpaullone intra-plate control as previously described (21). Assay effectiveness including S:N (a signal to noise ratio) and Z’ (an index of assay quality (37)) were calculated for both parameters using the same respective control. Both parameters were normalized to the average signal of DMSO (vehicle)-treated wells as a zero percent neutral control and the average signal of 57.5 µM kenpaullone-treated wells as maximum PFK1-mEGFP assemblies. Cell counts based on the number of nuclei were used as a parameter of cytotoxicity, which was also normalized to the average signal of DMSO-treated wells as a zero percent neutral control and the average signal of 57.5 µM SU9516-treated wells as maximum PFK1-mEGFP assemblies. The respective normalized data from each assay plate was corrected using DMSO only treated assay plates at the beginning and end of each library screen. In-house software (http://ncgc.nih.gov/pub/openhts/curvefit/) was used to fit the resulting intra-plate titration data to the standard hill equation, and concentration-response curves were classified by a titration curve as previously described (38). Curve fits assignments of class 1.1, 1.2, 2.1 and 2.2s were considered active (i.e., promotion of the PFK1-mEGFP assemblies) and visually confirmed. These classes included compounds which induced a full titration, a partial titration response, or a bell-shaped curve when the formation of PFK1-mEGFP assemblies was plotted versus log compound concentration. These data for both PFK1 assembly parameters and cell count cytotoxicity were refit in GraphPad Prism (GraphPad Software, Inc.) with nonlinear regression log(compound) vs. response-variable slope (four parameters) fit. Bell-shaped curves were fit in GraphPad Prism with nonlinear regression and the equation of:

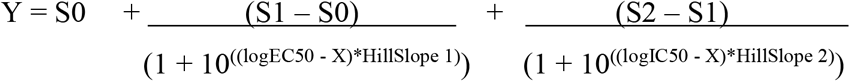

EC_50_ values were also calculated using GraphPad Prism (GraphPad Software, Inc) using non-linear four parameter curve fitting. In addition, images obtained from wells treated with compounds resulting in active curve classes for PFK1 assembly parameters were visually inspected in the GE workstation software to confirm the impact of the compounds. Compounds which had an active curve class but were categorized as cytotoxic by cell count were not considered.

### Cell cycle analysis

HeLa-T-PFK1G cells were plated in 6-well plates for a confluency of 70-90% in RPMI 1640 supplemented with 10% dialyzed FBS in the presence of 1 µg/mL doxycycline. The following day, cells were treated with SU9516 or DMSO for 24 hours. Cells were then harvested and analyzed for cell cycle progression as previously described (39). Briefly, hundreds of thousands of cells were removed from 6-well plates using trypsin, washed with 1x PBS (pH 7.4), resuspended in ice-cold PBS and fixed with ice-cold methanol for 20 minutes at 4° C. Cells were then washed and resuspended in the cell cycle buffer, which contains 30 µg/mL propidium iodide and 100 µg/mL RNase A (Thermo Cat # EN0531) in 1x PBS (pH 7.4). After being incubated for at least 45 minutes, flow cytometry was performed with a CyAn ADP (Beckman Coulter) running Summit V 4.00, equipped with a 488 nm laser line and 530/40 and 613/20 emission filters. Using FlowJo (FlowJo, LLC), cell populations were gated based on the expression of PFK1-mEGFP in cells as well as their size and granularity. The cell cycle progression was then assessed by the Dean-Jett-Fox analysis using a Cell Cycle platform available in FlowJo (40).

### Small-hairpin RNA (shRNA) interference

A shRNA expression system of the GIPZ Lentiviral shRNA Transfection Starter Kit (Dharmacon Inc.) containing V2LHS-153607 (Cat# RHS4430-200180936, GIPZ-shAURKA#1, TAAATAACACTTTAAGAGC), V2LHS-12364 (Cat# RHS4430-200216217, GIPZ-shAURKA#2, TATAAGTAGCAGTTCTCTG), V3LHS-638501 (Cat# RHS4430-200224041, GIPZ-shAURKA#3, CTATGAATAACTCTCTTCG) and validated non-silencing control (GIPZ Lentiviral Non-Silencing shRNA Control, shControl) was used according to the manufacturer’s protocol. Each shRNA plasmid was transfected using Lipofectamine 2000 as described above. Only transfected cells expressing an internal marker, turboGFP, of the shRNA expression system were subjected for fluorescence live-cell microscopy.

### Western blot analysis

HeLa cells were cultured in RPMI 1640 supplemented with 10% dialyzed FBS, transfected with each shRNA, and harvested after washing twice with 5 mL of 1x PBS. Harvested cells were stored at -80 ^o^C overnight, thawed in ice next day, and lysed in the RIPA buffer (20 mM Tris/HCl (pH 7.5), 140 mM NaCl, 1% Triton X-100, 0.5 % sodium deoxycholate, 0.1 % sodium dodecyl sulfate (SDS), and 10 % glycerol) that contains protease inhibitors (Pierce, Cat# 88666) and phosphatase inhibitors (Pierce, Cat# 88667). Protein concentrations of cell lysates were determined using Bradford protein assay (BioRad). We then prepared 10% SDS-PAGE gels and loaded 60 µg of each sample per lane. BioRad Semi Dry Transfer System was used to transfer resolved proteins to a PVDF membrane. The membrane was soaked in 5 % non-fat milk that was prepared in 1x TBST (10 mM Tris (pH 7.5), 150 mM NaCl, and 0.5% Tween 20), followed by washing three times with 1x TBST. Then, a primary antibody of rabbit anti-Aurora Kinase A (D3E4Q) (1:1000, Cell Signaling Technology, Cat#14475) was incubated with the membrane overnight at 4 °C. Subsequently, a secondary antibody of donkey anti-rabbit-AlexaFluor790 (1:10,000, Jackson Immunoresearch, Cat# 711-655-152) was incubated with the membrane for 1-2 hours at room temperature. LI-COR Odyssey Sa Near-IR imager was then used to image the membrane, and ImageStudioLite (LI-COR) was used for quantitative analysis.

## Results

### Development of a cell-based assay reporting the formation of PFK1-mEGFP assembly

To develop a cell-based assay for small-molecule library screening, we first adapted the propensity of PFK1-mEGFP to form spatial assemblies in various cancer cell lines (4) as an intracellular reporter. To begin with, as reported previously with Hs578T cells (4), we evaluated what percentage of transiently transfected HeLa cells with PFK1-mEGFP showed the formation of PFK1 assemblies. We found that 51 ± 5 %, 22 ± 3 % and 23 ± 2 % of HeLa cells with PFK1-mEGFP exhibited small- (i.e., < 0.1 µm^2^), medium- (i.e., < 3 µm^2^) and large-sized (i.e., < 8 µm^2^) assemblies, respectively, while 5 ± 2% of HeLa cells showed no assembly (**Figure 1A**). However, due to various expression levels of PFK1-mEGFP (4) and size distribution of its assemblies at single-cell levels among transiently transfected HeLa cells (**Figure 1A**), we decided to establish an inducible expression system using HeLa Tet-On 3G cells (Clontech), in which PFK1-mEGFP was stably expressed by addition of doxycycline (1 µg/mL). This stably transfected cell line, namely the ‘HeLa-T-PFK1G’ cells, was then used in this work to identify small molecules that promoted the formation of PFK1-mEGFP assemblies.

**Figure 1.**
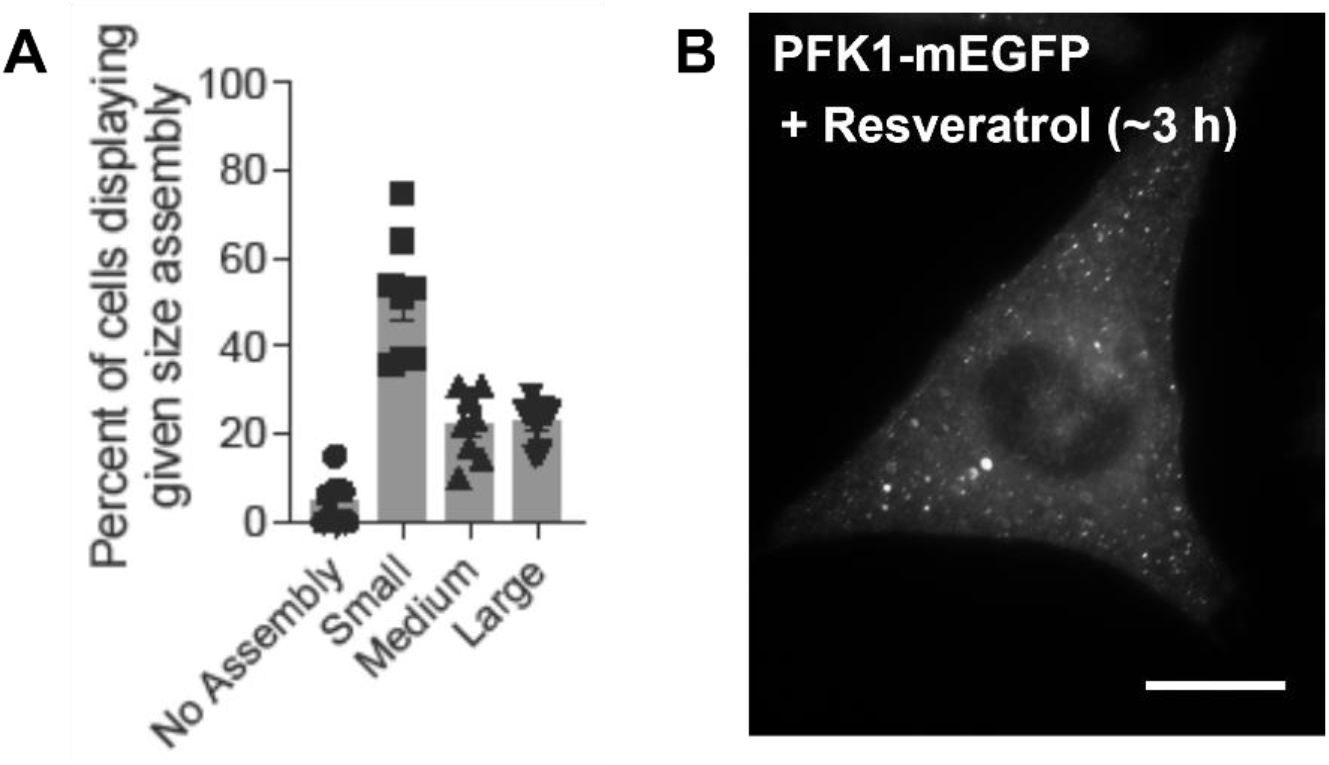
PFK1-mEGFP assemblies in HeLa cells. (**A**) Analysis of the formation of PFK1-mEGFP assemblies from transiently transfected HeLa cells, where small-sized assemblies are defined to have < 0.1 µm^2^, medium < 3 µm^2^, and large < 8 µm^2^. Error bars represent standard errors of the means of at least three independent trials, average of 1165 cells. (**B**) Resveratrol-induced formation of PFK1-mEGFP assemblies in fixed HeLa-T-PFK1G cells. A representative image was selected among at least 1000 cells. Scale bar, 10 µm.

Development of a cell-based qHTS assay also required a positive control compound to ensure the assay’s robustness and reproducibility. Due to the reported implication of resveratrol, a polyphenolic compound, on glycolysis (41,42), we first tested transiently transfected HeLa cells that expressed PFK1-mEGFP with resveratrol (200-250 µM) and observed that ∼80-90 % of HeLa cells exhibited medium- and large-sized assemblies in their cytoplasm. We then confirmed the effect of resveratrol on the formation of PFK1 assemblies in the stably transfected HeLa-T-PFK1G cells after fixation (**Figure 1B**). Although a specific mechanism of action of resveratrol is not clear yet due to its numerous biological implications (43), as seen for flavonoids in general (44,45), the observed effect of resveratrol on the formation of PFK1 assembly was sufficient enough to serve as a positive control in high-throughput assays. In short, as described in the Methods section, we successfully worked on adapting our microscopy-based cell assay into a high-throughput compatible assay with computer-aided high-content analysis.

### Identification of SU9516 from cell-based qHTS with the LOPAC library

We then performed the cell-based qHTS assay using the LOPAC library, which contains 1280 pharmacologically active compounds. Analysis of two independent screens of the LOPAC library initially identified 16 active compounds (1.25 % of the library) (**Supplementary Table 1**). A validation screen was subsequently performed testing these 16 compounds in a 10-point 1:3 titration, ranging from 57.5 µM to 8.7 nM. Cell images collected from wells treated with 6.4 µM or higher concentrations of the compounds were manually inspected for PFK1-mEGFP assemblies to evaluate the robustness of our semi-automated computer-aided high-content analysis algorithm. As a result, 8 compounds (0.63% of the LOPAC library) appeared to promote the formation of PFK1-mEGFP assemblies in HeLa-T-PFK1G cells with minimal cytotoxicity based on nuclei count at their active concentrations (AC_50_) (**Table 1**).

**Table 1.**
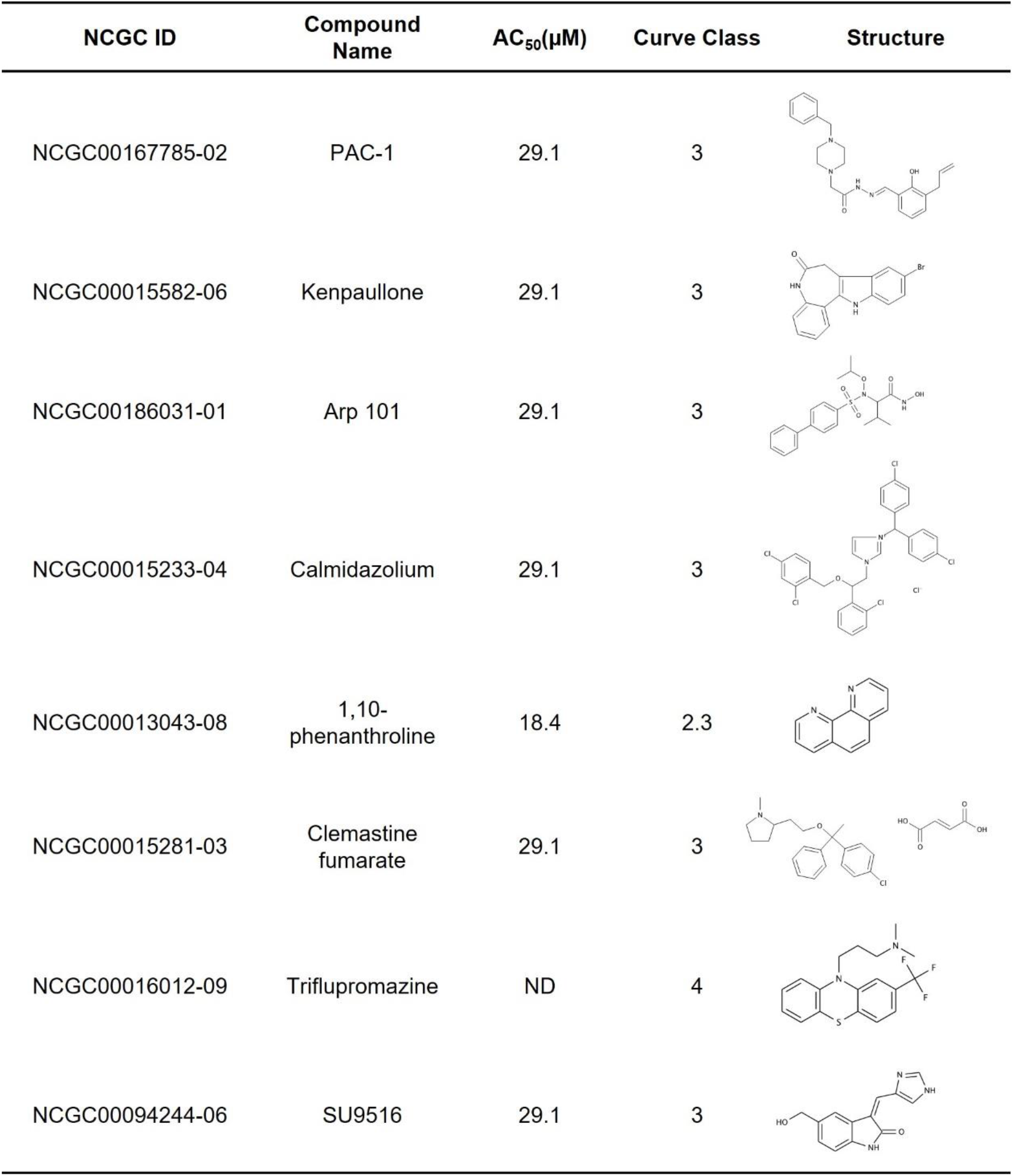
Hit compounds from the LOPAC screen.

| NCGC ID | Compound Name | AC<sub>50</sub>(μM) | Curve Class | Structure |
| --- | --- | --- | --- | --- |
| NCGC00167785-02 | PAC-1 | 29.1 | 3 |  |
| NCGC00015582-06 | Kenpaullone | 29.1 | 3 |  |
| NCGC00186031-01 | Arp 101 | 29.1 | 3 |  |
| NCGC00015233-04 | Calmidazolium | 29.1 | 3 |  |
| NCGC00013043-08 | 1,10-phenanthroline | 18.4 | 2.3 |  |
| NCGC00015281-03 | Clemastine fumarate | 29.1 | 3 |  |
| NCGC00016012-09 | Triflupromazine | ND | 4 |  |
| NCGC00094244-06 | SU9516 | 29.1 | 3 |  |

These 8 compounds were further assessed for their ability to induce PFK1-mEGFP assemblies in HeLa-T-PFK1G cells under high-resolution fluorescence single-cell microscopy. When we treated cells with 57.5 µM and 10 µM of each compound for 5 hours (**Figure 2A and 2C**) and 25 hours (**Figure 2B and 2D**), respectively, only SU9516 (NCGC00094244-06) was capable of inducing PFK1-mEGFP assemblies robustly in HeLa-T-PFK1G cells. In detail, SU9516 induced medium- and large-sized PFK1 assemblies in 27 ± 5 % and 40 ± 5% of the cells by the treatment of 57.5 µM for 5 hours and the treatment of 10 µM for 25 hours, respectively (**Figure 2**). Furthermore, an effective concentration (EC_50_) of 6.8 µM SU9516 was determined by titrating various concentrations of SU9516 with 25 hours incubation under high-resolution imaging (**Supplementary Figure 1**). Alternatively, we also transiently transfected PFK1-mEGFP into parental HeLa cells and treated with 57.5 µM of SU9516. After 5 hours, we were able to monitor the promotion of medium- and large-sized assemblies at single-cell levels (**Supplementary Figure 2**). Taken together, we confirmed that SU9516 was capable of promoting PFK1-mEGFP assemblies in HeLa cells.

**Figure 2.**
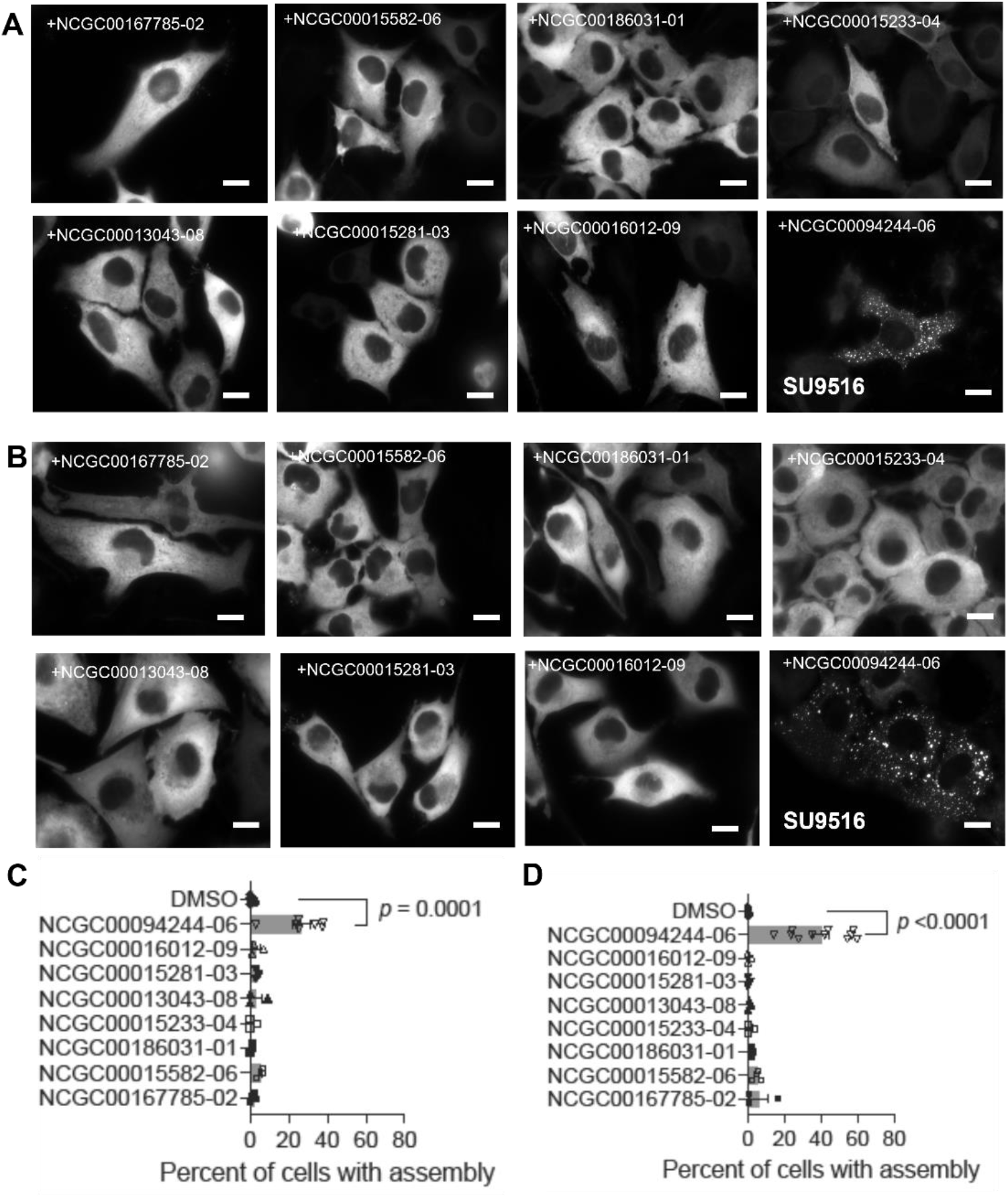
High-resolution imaging validation of LOPAC hit compounds. (**A** and **B**) HeLa-T-PFK1G cells were treated with 8 hit compounds (Table 1) at 57.5 µM for 5 hours (**A**) or 10 µM for 25 hours (**B**) and assessed for their impact on the formation of PFK1-mEGFP assemblies. Representative images were selected from at least three independent experiments. (**C** and **D**) Quantitation of the average percentage of HeLa-T-PFK1G cells showing PFK1-mEGFP assemblies after treatment with an indicated compound: 57.5 µM for 5 hours (**C**) or 10 µM for 25 hours (**D**). Error bars represent standard errors of the means of at least three independent trials. Statistical significance was determined using student’s two-tailed *t* test, at least 300 cells analyzed per condition. Scale bar, 10 µm.

Nevertheless, it is important to emphasize that multi-step imaging-based screening strategies are important to ensure the formation of PFK1-mEGFP assemblies at subcellular levels. Under high-resolution imaging, 7 compounds other than SU9516 did not induce PFK1-mEGFP assemblies at either concentration tested (**Figure 2**). False positive results from qHTS assays could be in part because non-specific phenotypic changes or aggregation of PFK1-mEGFP by small molecule treatment and/or cytotoxic stress could be miss-classified as PFK1-mEGFP forms assemblies at low resolution imaging platforms. Also, some compounds are known to produce artifactual fluorescence puncta during cell-based qHTS assays (24). When fluorescent signals were significantly measurable in non-PFK1-mEGFP expressing cells, or when fluorescent puncta response was greater in non-induced over doxycycline-induced PFK1-mEGFP expressing cells, we considered them as non-relevant to PFK1 assemblies. In this work, we found that kenpaullone, along with BIM II and BIM X as artifact-causing positive controls (24), produced non-relevant fluorescent puncta in HeLa cells (**Supplementary Figure 3**). However, formation of such fluorescent artifacts was negligible with SU9516 in our conditions (**Supplementary Figure 3A, 3E, and 3F**), supporting the authenticity of PFK1 assemblies induced by SU9516 (**Figure 3**). At the same time, relatively weak activities of compounds and thus their poor curve fits classification from qHTS assays might be also anticipated because of differences in optical resolution and magnification between two imaging platforms: the qHTS image analysis relied on cell images obtained by a 20x objective lens that has 0.45 numerical aperture while high-resolution fluorescence microscopy was performed with a 60x objective lens that has 1.45 numerical aperture. Collectively, SU9516 was the only compound from the LOPAC library (i.e., 0.078% of the library) that promoted PFK1-mEGFP assemblies in HeLa cells.

**Figure 3.**
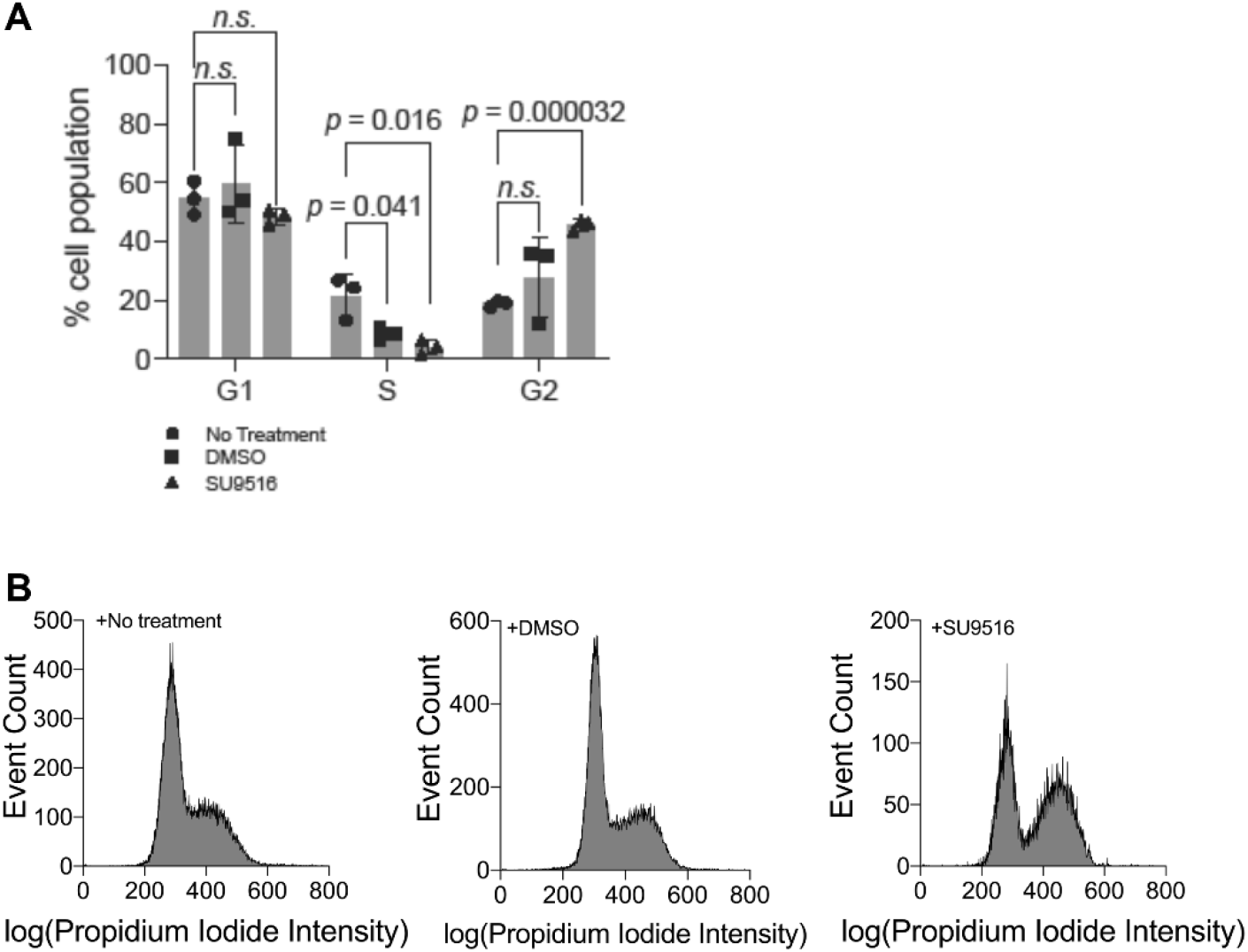
Effect of SU9516 on cell cycle progression of HeLa-T-PFK1G cells. Cells were treated with no small molecule, DMSO or 10 µM SU9516, and incubated for 24 hours. Then, cells were assessed for cell cycle progression by propidium iodide staining. (**A**) Changes of the average percentage of cells in each phase of a cell cycle following incubation with an indicated compound. Error bars represent standard deviations of at least three trials and statistical significance was determined using student’s two-tailed *t* tests, data represent at least 3000 cells analyzed per run. (**B**) Representative histograms of cell cycle progression of HeLa-T-PFK1G cells receiving either no treatment, DMSO, or 10 µM SU9516 for 24 hours. Histograms were selected from at least three independent experiments.

### Effect of SU9516 on cell cycle progression of HeLa cells

SU9516 has been reported to inhibit CDK2, thus impacting cell cycle progression (46-48). We performed flow cytometric analysis to investigate whether the treatment of SU9516 played a role in cell cycle progression of HeLa cells in our conditions. When we compared the distribution of cells at each phase of a cell cycle in the presence and absence of SU9516, we found that the population of HeLa cells in the S phase was significantly decreased from 8.7 ± 2.2 % to 4.2 ± 2.4% in the presence of SU9516, while the population of HeLa cells in the G2 phase was significantly increased from 27.8 ± 13.3 % to 45.5 ± 1.9 % (**Figure 3**). No significant change was observed in the population of HeLa cells in the G1 phase (i.e., 59.7 ± 13.2 % vs. 48.4 ± 2.7 %) (**Figure 3**). This result is indeed consistent with previous studies that showed the impact of SU9516 on a cell cycle of colon cancer cells (46). Therefore, the promotion of PFK1-mEGFP assemblies by SU9516 in HeLa cells appears to associate with the inhibition of CDK2 and thus cell cycle progression.

### Identification of SL-0101-1 and PF-03814735 from a kinase inhibitor-enriched library (KIEL) screening

Although SU9516 has an established molecular target of CDK2 (IC_50_ = 0.022 µM), mechanistically it is an ATP-competitive inhibitor (46). Many off-targets of SU9516, including but not limited to CDK1 (IC_50_ = 0.04 µM) and CDK4 (IC_50_ = 0.2 µM), have been already identified (49). To understand the primary target specificity or off-target effects of SU9516 on the promotion of PFK1 assemblies from our conditions, we further screened 2958 kinase inhibitors from a KIEL library that we prepared combining three smaller libraries: the Protein Kinase Inhibitor set (32,33), the Roche Kinase Inhibitor Library (34) and NCATS mechanism interrogation plates (35). Initially, nearly 8% of the library (i.e., 233 molecules) yielded qHTS curve fits of class 1 to class 2 based on an analysis of concentration-response curves (**Table 2**). However, the initial hit analysis seemed to correlate with high cytotoxicity for most active compounds. Accordingly, compounds demonstrating cytotoxicity based on nuclei count or visually confirmed to be experimental artifacts due to, for example, precipitation of compounds in growth medium, were eliminated from further consideration, thereby narrowing down the number of active molecules to 24; among them, 8 compounds belonged to a curve class 1 while 16 compounds were in a curve class 2 (**Table 3**). One step further, of the 24 active compounds from the KIEL library (**Table 3**), 5 compounds were reported to primarily inhibit known off-targets of SU9516, including Aurora kinases, CDKs, and RSK (**Table 4**) (49-54). Therefore, the 5 focused compounds from the KIEL library were subjected to further validation assays under high-resolution fluorescence single-cell imaging.

**Table 2.**
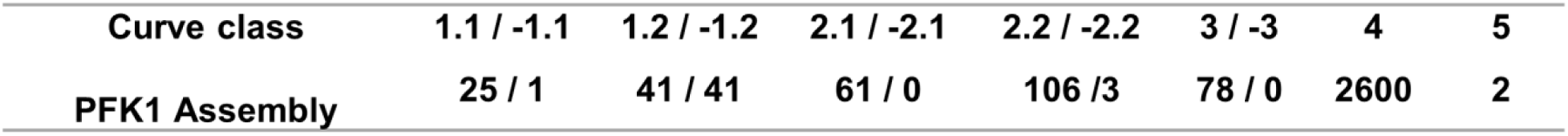
Curve class distribution of small molecules from the kinase inhibitor-enriched library screen.

**Table 3.** Small molecules that induced PFK1-mEGFP assemblies from the kinase inhibitor-enriched library.

| NCGC ID | Curve Class | AC50 (μM) | Primary Target | Reference |
| --- | --- | --- | --- | --- |
| NCGC00263213-01 | 1.1 | 7.7 | IKK-2 (IKK-beta) Inhibitor | Sommers et al J Pharmacol Exp Ther 330 377-88 (2009) |
| NCGC00346518-01 | 1.2 | 2.7 | Carbapenem Antibiotic | Pryka Ann Pharmacother 28 1045-54 (1994) |
| NCGC00348107-01 | 1.2 | 9.7 | CSF1R (c-FMS) Inhibitor | USA Patent US7705042 |
| NCGC00094087-06 | 1.4 | 9.7 | Inosine 5'-Monophosphate Dehydrogenase (IMPDH) Inhibitors | Koyama et al Biochem Pharmacol 32 3547-53 (1983) |
| NCGC00159346-05 | 1.4 | 9.7 | Fungal Squalene Monooxygenase Inhibitor | Ryder Clin Exp Dermatol 14 98-100 (1989) |
| NCGC00344512-01 | 1.4 | 9.7 | Opioid receptor antagonist | Schmidhammer et al J Med Chem 32 418-21 (1989) |
| NCGC00167513-03 | 1.4 | 1.2 | EGFR (HER1; erbB1) Inhibitor | Hennequin et al J Med Chem 45 1300-12 (2002) |
| NCGC00165811-03 | 1.4 | 48.5 | Inhibitor of nuclear factor kappa b kinase subunit beta (IKK-2) | Onai et al Cardiovasc Res 63 51-9 (2004) |
| NCGC00346698-01 | 2.1 | 19.3 | mTORC1/2 inhibitor | Pike et al Bioorg Med Chem Lett 23 1212-6 (2013) |
| NCGC00346747-02 | 2.1 | 48.5 | NFkappaB-inducing kinase Inhibitor | USA Patent US2011086834 |
| NCGC00347280-01 | 2.1 | 7.7 | IKK-2 (IKK-beta) Inhibitor | Murata et al Bioorg Med Chem Lett 14 4019-22 (2004) |
| NCGC00241982-03 | 2.1 | 30.6 | ROCK 1, ROCK 2 Inhibitor | Stavenger et al J Med Chem 50 2-5 (2007) |
| NCGC00346553-01 | 2.1 | 24.3 | CDK7 | Ali et al Cancer Res 69 6208-15(2009) |
| NCGC00263129-01 | 2.1 | 8.6 | CDK4, 6 | Fry et al Mol Cancer Ther 3 1427-38 (2004) |
| NCGC00018248-08 | 2.1 | 17.2 | Cyclooxygenase-1/2 Inhibitor | Noble et al Drugs 51 424-30 (1996) |
| NCGC00166111-04 | 2.1 | 48.5 | Estrogen Receptor (ER) Agonist (nuclear) | Clark et al J Anim Sci 49 46-65 (1979) |
| NCGC00181306-03 | 2.1 | 48.5 | Tubulin depolymerization inhibitor | Plenta Semin Oncol 28 3-7 (2001) |
| NCGC00346893-01 | 2.1 | 30.6 | MAPKAP-K1 (RSK; p90Rsk) Inhibitor | Smith et al Cancer Res 65 1027-34 (2005) |
| NCGC00346542-01 | 2.1 | 48.5 | Aurora kinase inhibitor | Hauf et al Journal of Cell Biology 161 281-94 (2003) |
| NCGC00346652-01 | 2.1 | 43.2 | Aurora kinase inhibitor | Jani et al Mol Cancer Ther 9 883-94 (2010) |
| NCGC00263125-01 | 2.2 | 48.5 | mTORC1/2 inhibitor | Weinberg Anticancer Drugs 27 475-87 (2016) |
| NCGC00346681-01 | 2.3 | 9.7 | PI3K Inhibitor | Chang et al Clin Cancer Res 15 7116-26 (2011) |
| NCGC00346959-01 | 2.3 | 38.5 | Lck Kinase Inhibitors | Martin et al J Med Chem 49 4981-91 (2006) |
| NCGC00015420-09 | 2.4 | 48.5 | Casein Kinase II (CK2) Inhibitor | Srinivas et al Med Res Rev 27 591-608 (2007) |

**Table 4.**
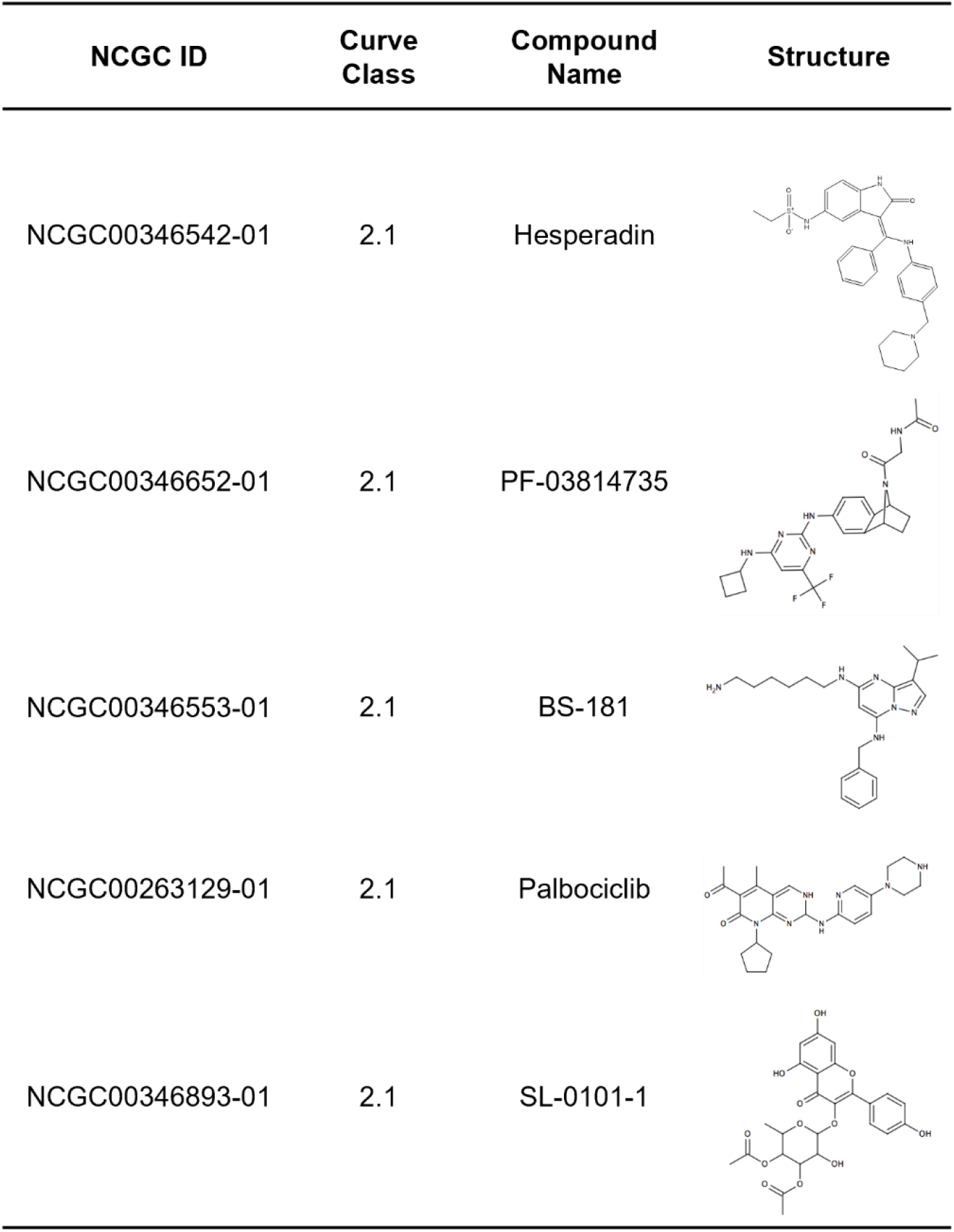
Small molecules from the kinase inhibitor-enriched library screen which inhibit off-targets of SU9516.

| NCGC ID | Curve Class | Compound Name | Structure |
| --- | --- | --- | --- |
| NCGC00346542-01 | 2.1 | Hesperadin |  |
| NCGC00346652-01 | 2.1 | PF-03814735 |  |
| NCGC00346553-01 | 2.1 | BS-181 |  |
| NCGC00263129-01 | 2.1 | Palbociclib |  |
| NCGC00346893-01 | 2.1 | SL-0101-1 |  |

Of the 5 compounds, 2 molecules were confirmed to significantly induce PFK1-mEGFP assemblies in HeLa-T-PFK1G cells (**Figure 4**). Briefly, we found that 5-10 µM PF-03814735 (NCGC00346652-01, a pan inhibitor of Aurora kinases A and B) showed the most significant impact on the formation of PFK1 assemblies in ∼95% of HeLa-T-PFK1G cells after 1-2 hours treatment. More than 90% of the assembly-positive cells exhibited medium-sized assemblies in their cytoplasm. 10 µM SL-0101-1 (NCGC00346893-01, RSK inhibitor) also strongly promoted PFK1 assemblies in ∼77% HeLa-T-PFK1G cells after 4-5 hours incubation. Most of the assembly-positive cells (i.e., > 90%) again displayed medium-sized assemblies. We then confirmed that artifactual fluorescent puncta by SL-0101-1 and PF-03814735 were non-relevant or negligible, unlike kenpaullone and BIM compounds (**Supplementary Figure 3**), when tested in HeLa-T-PFK1G cells without doxycycline as well as parental HeLa cells (**Supplementary Figure 4**). However, the other 3 molecules; inhibitors of CDK7 (NCGC00346553-01, BS-181, 20 µM), CDK4/6 (NCGC00263129-24, palbociclib, 20 µM) and Aurora kinase B (NCGC00346542-1, hesperidin, 10-50 µM), did not promote PFK1-mEGFP assemblies in the cytoplasm of HeLa-T-PFK1G cells. Since the inhibition of Aurora kinase B by hesperadin (**Figure 4E-F**) did not trigger the formation of cytoplasmic assemblies of PFK1, the profound positive activity of PF-03814735 on the formation of PFK1 assembly (**Figure 4G-H**) appeared to be due to the A isoform-specific inhibition of Aurora kinase. Taken all together, our qHTS assays identified three active compounds (SU9516, SL-0101-1, and PF-03814735) that have been characterized to inhibit each node of the RSK-CDK2-AURKA signaling cascade, respectively, thereby revealing the importance of the kinase cascade in regulation of PFK1 assemblies in human cells.

**Figure 4.**
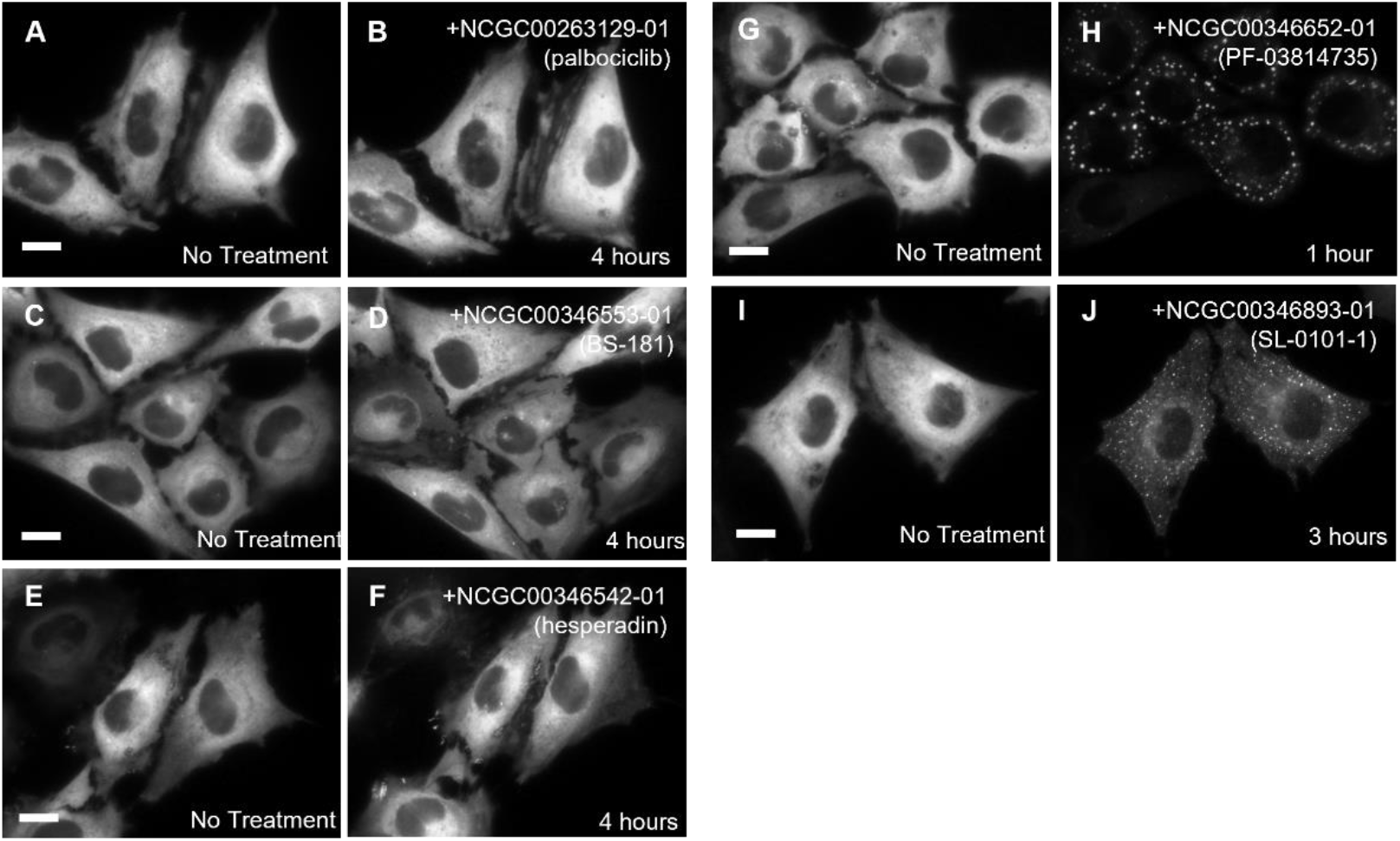
High-resolution imaging validation of the kinase inhibitor-enriched library (KIEL) compounds. HeLa-T-PFK1G cells were treated with 5 hit compounds from the KIEL library (Table 4). Treatment of 20 µM palbociclib for 4 hours (**A-B**), 20 µM BS-181 for 4 hours (**C-D**) or 50 µM hesperadin (**E-F**) for 4 hours barely promoted PFK1-mEGFP assembly. However, treatment of 5 µM PF-03814735 for 1 hour (**G-H**) or 10 µM SL-0101-1 for 2-3 hours (**I-J**) significantly promoted the formation of PFK1-mEGFP assemblies in HeLa-T-PFK1G cells. Representative images were selected from at least three independent experiments. Scale bar, 10 µm.

### Promotion of PFK1 assemblies by a signaling cascade activating Aurora kinase A (AURKA)

It appears that the cascade of three protein kinases we identified in this work forms a specific signaling axis in human cells. We hypothesized that AURKA might be the ultimate target of all the three identified inhibitors to regulate PFK1 assemblies in HeLa cells. We knocked down AURKA using shRNAs to validate our pharmacological molecule-based studies. First, western blot analysis showed that among three shRNAs we tested against AURKA (i.e., shAURKA1/2/3), shAURKA2 robustly showed ∼56% knockdown efficiency relative to shControl using scrambled shRNA sequence (**Figure 5A-B**). Next, to evaluate the impact of shAURKA2 on the formation of PFK1 assemblies at single-cell levels, we carried out high-resolution fluorescence single-cell microscopy with dually transfected HeLa cells expressing both PFK1-mCherry and shAURKA2. When we analyzed the formation of PFK1 assemblies in the presence of shAURKA2 over a negative control (shControl), we observed that knockdown of AURKA decreased ∼48 % of HeLa cells showing small-sized PFK1 assemblies but increased ∼33 % and ∼48 % of HeLa cells displaying medium- and large-sized PFK1 assemblies, respectively (**Figure 5C-F**). Collectively, this study reveals that the activity of AURKA is important for PFK1-mediated glucosome assemblies, particularly for the formation of medium- and large-sized glucosomes.

**Figure 5.**
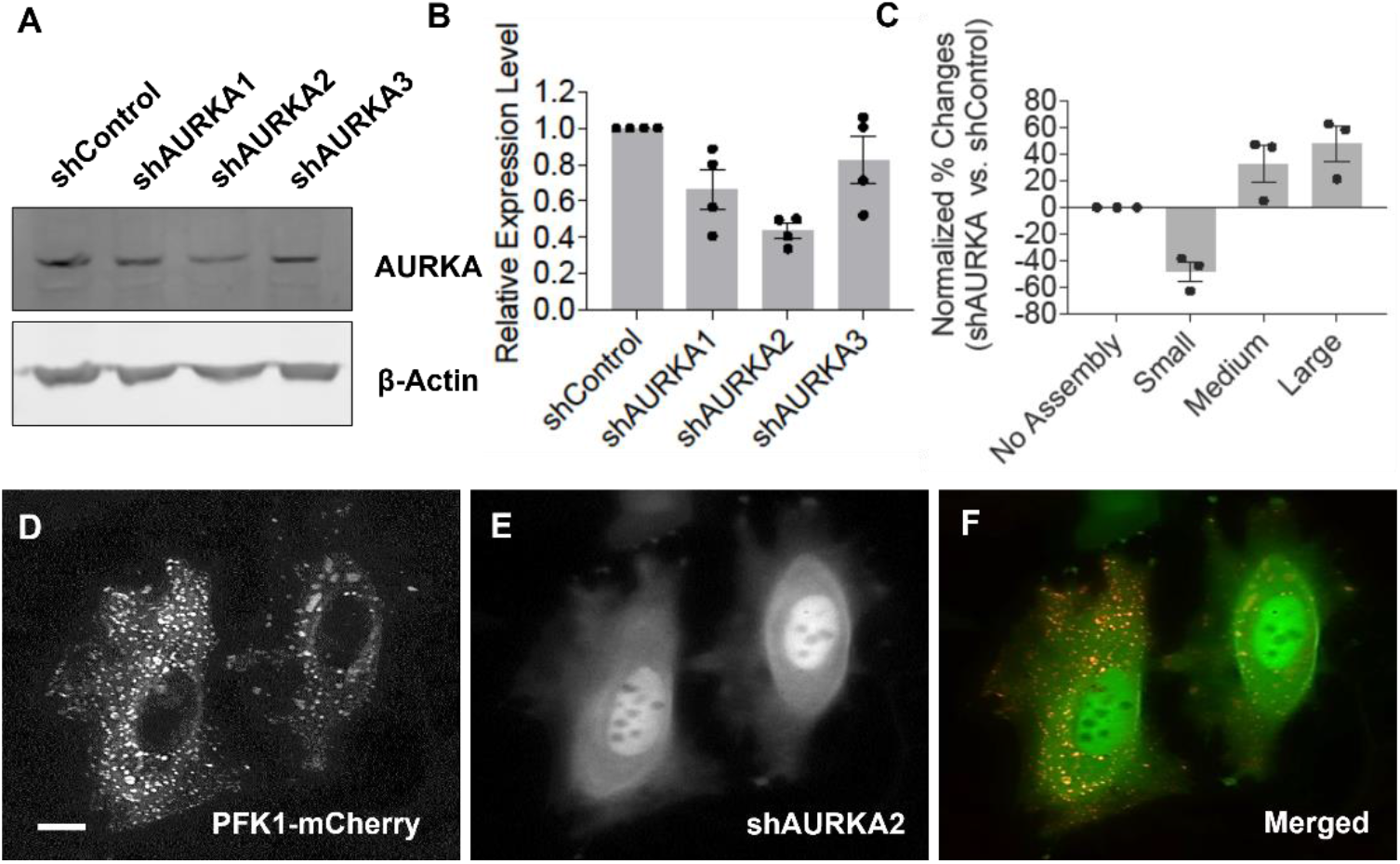
Knockdown of Aurora kinase A (AURKA). HeLa cells were transfected with three shRNAs targeting AURKA, respectively. (**A-B**) Western blot analysis showed that shAURKA2 robustly reduced the expression level of AURKA in HeLa cells. Error bars represent standard deviations from at least four independent trials. (**C**) Fluorescence live-cell imaging revealed that the AURKA knockdown by shAURKA2 decreased the population (%) of transiently transfected HeLa cells showing small-sized assemblies (Small, -48.4 %) while increased the populations (%) of HeLa cells displaying medium- and large-sized assemblies (Medium, +32.5 %; Large, +47.6 %). A least three independent imaging experiments were carried out to graph the averaged effects of shAURKA2 vs. shControl on each size of PFK1-mCherry assembly. (**D-F**) Representative images showing glucosomes (PFK1-mCherry in the cytoplasm of HeLa cells, **D**) in the presence of shAURKA2 (shown as internal turboGFP control signals in the nucleus, **E**). Scale bar, 10 µm.

## Discussion

We have previously characterized a multienzyme metabolic assembly, the glucosome, that is involved in glucose metabolism in human cells (4). However, it has remained largely elusive how glucosomes are reversibly assembled and disassembled to functionally regulate glucose metabolism in human cells. In this work, we developed a cell-based qHTS assay by which the formation of PFK1 assemblies was monitored from stably transfected HeLa-T-PFK1G cells in response to a small molecule. By screening more than 4200 pharmacologically active compounds, we identified three small molecules, SL-0101-1, SU9516 and PF-03814735, that significantly promoted the formation of PFK1 assemblies in HeLa cells through the inhibition of the RSK-CDK2-AURKA signaling axis (**Figure 6**). Briefly, 95% of HeLa-T-PFK1G cells showed the promotion of PFK1 assemblies in 1-2 hours during the treatment of 5-10 µM PF-03814735 (AURKA inhibitor, **Figure 4**), 77% of the cells in 4-5 hours by 10 µM SL-0101-1 (RSK inhibitor, **Figure 4**), and 40% of the cells in 25 hours by 10 µM SU9516 (CDK2 inhibitor, **Figure 2**). Considering the population (%) of HeLa cells responding to each inhibitor and the treatment duration required for the cellular response, we propose that AURKA would be the ultimate target that all the three identified compounds are intended to regulate, to control PFK1 assemblies in HeLa cells. However, the off-target effect of SU9516 to CDK1 (IC_50_ = 0.04 µM) over its primary target CDK2 (IC_50_ = 0.022 µM) cannot be ignored (46) (**Figure 6**). Also, RSK and AURKA are shown to activate CDK1 through alternative pathways to regulate cell cycle progression (**Figure 6**). We cannot completely rule out the possibility of CDK1 involvement in regulation of PFK1 assembly in HeLa cells. Nevertheless, the RSK-CDK2-AURKA signaling axis and even CDK1-associated pathways play an important role during cell cycle progression (26-29). Considering that PFK1 is a scaffold enzyme for multienzyme glucosome assemblies (4,15,55), our results provide strong evidence that cell cycle-associated signaling cascades regulate reversible formation of PFK1-mediated glucosome assemblies in human cells.

**Figure 6.**
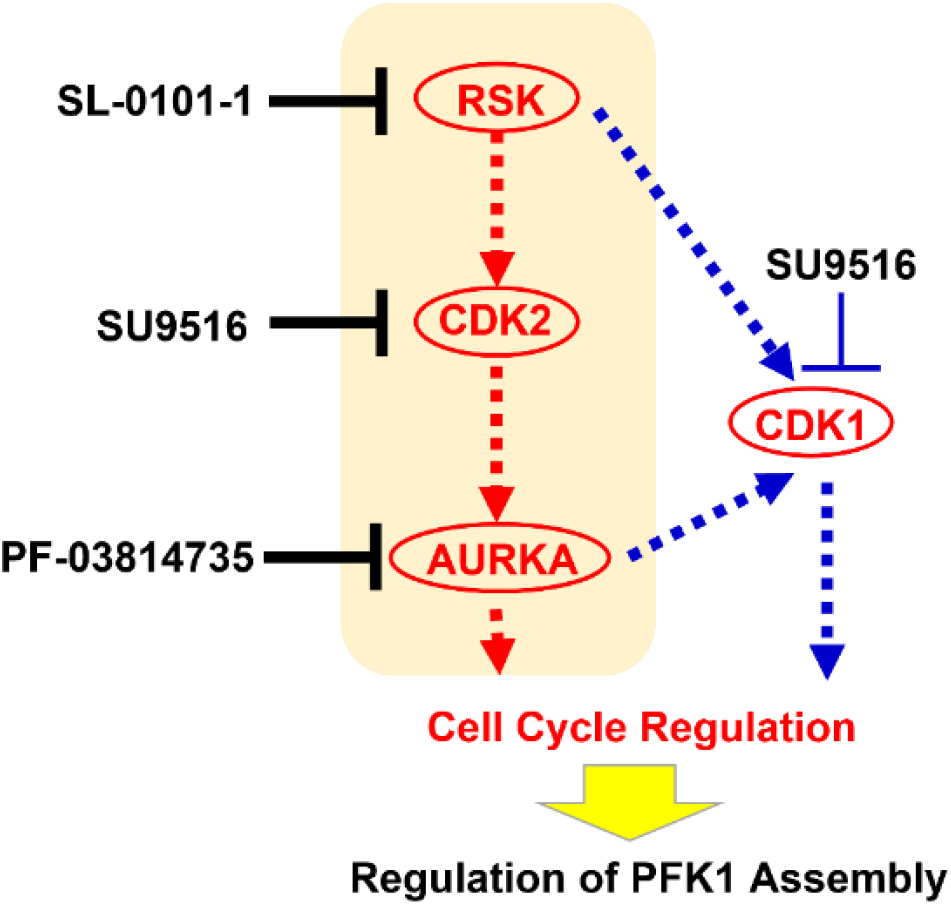
A schematic diagram of the identified cell cycle-associated signaling cascades that influence the promotion of PFK1-mediated glucosome assembly in HeLa cells. Briefly, ribosomal protein S6 kinase (RSK) downregulates a cyclin-dependent kinase inhibitor, p27^Kip1^, that inhibits cyclin-dependent kinase 2 (CDK2). In turn, CDK2 phosphorylates retinoblastoma-associated protein, Rb1, to activate Aurora kinase A (AURKA), followed by the promotion of G1 and G1/S transition. Alternatively, RSK and AURKA are shown to activate CDK1 through alternative pathways to regulate G2/M transition.

At the same time, all the identified protein kinases, RSK, CDK2 and AURKA, have been also demonstrated to directly influence glycolysis in human cancer cells. For instance, RSK directly phosphorylates the regulator domain of PFKFB2, thus increasing its catalytic activity to control glycolysis in melanoma cells (56). CDK2 is also found to decrease the activity of triosephosphate isomerase by direct phosphorylation in HeLa cells (57). Knockdown of hexokinase II stalled cancer cells in the G1 phase through the inhibition of CDK2 (11). AURKA is known to directly interact with and phosphorylate lactate dehydrogenase, thus increasing glycolysis and lactate production in cancer cells (58). Therefore, it is possible that the RSK-CDK2-AURKA signaling cascade is capable of directly controlling glycolytic flux through the regulation of glucosome assemblies to meet metabolic demands during a cell cycle of human cancer cells.

Collectively, many metabolic enzymes have been found to be reversibly compartmentalized into condensates, rings, rods, and/or filaments as membraneless structures in living cells (1). Although phase separation-mediated formation of multienzyme condensates has been recently demonstrated in yeast and human cells (5,6), underlying mechanisms that drive such various enzyme compartments in cells are largely unknown. We demonstrate here that qHTS approaches would be advantageous in identifying heretofore unidentified cellular mechanisms that regulate compartmentalization of metabolic enzymes in living cells, thus offering novel insights for therapeutic intervention.

## Acknowledgement

We would like to thank Dr. S. Ostrand-Rosenberg (UMBC), Dr. G. Szeto (UMBC), M. Zhang for assisting our data acquisition by flow cytometry, M. Jeon for critically reading the manuscript, and L. Harris for our literature mining, and Dr. R. MacArthur for informatics support (NCATS). We also thank Drs. D. Drewry and W. Zuercher (University of North Carolina, Chapel Hill) for the PKIS and Dr. P. Gillespie for the Roche Kinase Inhibitor Library.

## Funding Discloser

AACR-Bayer Innovation and Discovery Grant (16-80-44-ANSO; S.A)

NIH/NIGMS R01GM125981 (S.A)

NIH/NCI R03CA219609 (S.A)

NIH/NIGMS R01GM134086 (M.K.)

NIH/NIGMS T32GM066706 (D.L.S. and C.L.K.)

NIH/NCATS 1ZIATR000052-04 (J.I.)

