## Supplemental for "High-content quantitative high-throughput screening identifies a cell cycle-associated signaling cascade that regulates a multienzyme metabolic assembly for glucose metabolism"

**Supplementary Table 1. Initial 16 hit compounds from the LOPAC screen.**

| NCGC ID | Compound Name |
|---|---|
| NCGC00013043-08 | 1,10-Phenanthroline monohydrate |
| NCGC00015856-07 | Prochlorperazine |
| NCGC00014483-11 | Nortriptyline hydrochloride |
| NCGC00015233-04 | Calmidazolium chloride |
| NCGC00015281-03 | Clemastine fumarate |
| NCGC00015376-06 | N,N-Dihexyl-2-(4-fluorophenyl)indole-3-acetamide |
| NCGC00016888-03 | Fluoxetine hydrochloride |
| NCGC00015582-06 | Kenpaullone |
| NCGC00016012-09 | Promazine hydrochloride |
| NCGC00016012-09 | Triflupromazine hydrochloride |
| NCGC00016888-03 | S-(+)-Fluoxetine hydrochloride |
| NCGC00094244-06 | SU 9516 |
| NCGC00167785-02 | PAC-1 |
| NCGC00186031-01 | ARP 101 |
| NCGC00015701-05 | DL-alpha-Methyl-p-tyrosine |
| NCGC00094144-06 | L-alpha-Methyl-p-tyrosine |

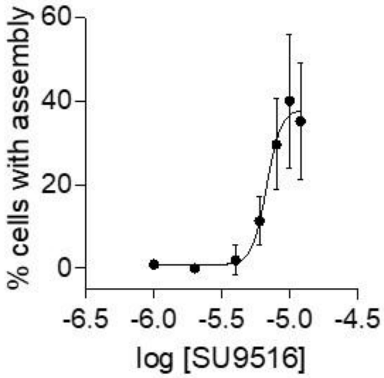


**Supplementary Figure 1. EC$_{50}$ measurement of SU9516.** HeLa-T-PFK1G cells were treated with SU9516 in titration for 25 hours and the number of cells showing PFK1 assemblies was assessed. Error bars represent standard deviations of at least three independent trials. At least 300 cells were analyzed per condition.

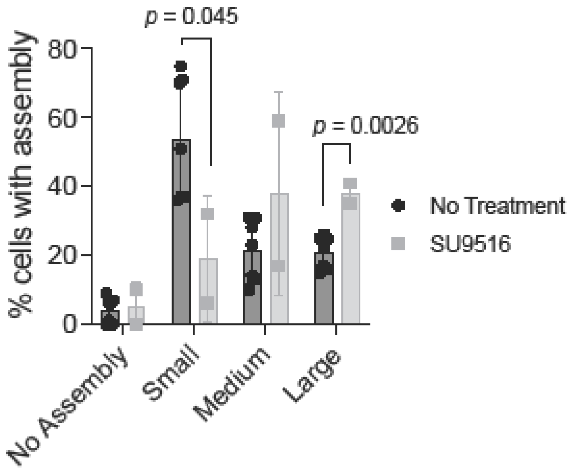


**Supplementary Figure 2**. **Effect of SU9516 on transiently expressed PFK1-mEGFP in HeLa cells.** HeLa cells transiently expressing PFK1-mEGFP were treated with 57.5 µM SU9516 for 5 hours. Following treatment, the percent of cell population containing no assembly, small, medium, or large sized assemblies were assessed. Error bars represent standard errors. Statistical significance was determined using student's two-tailed t test.

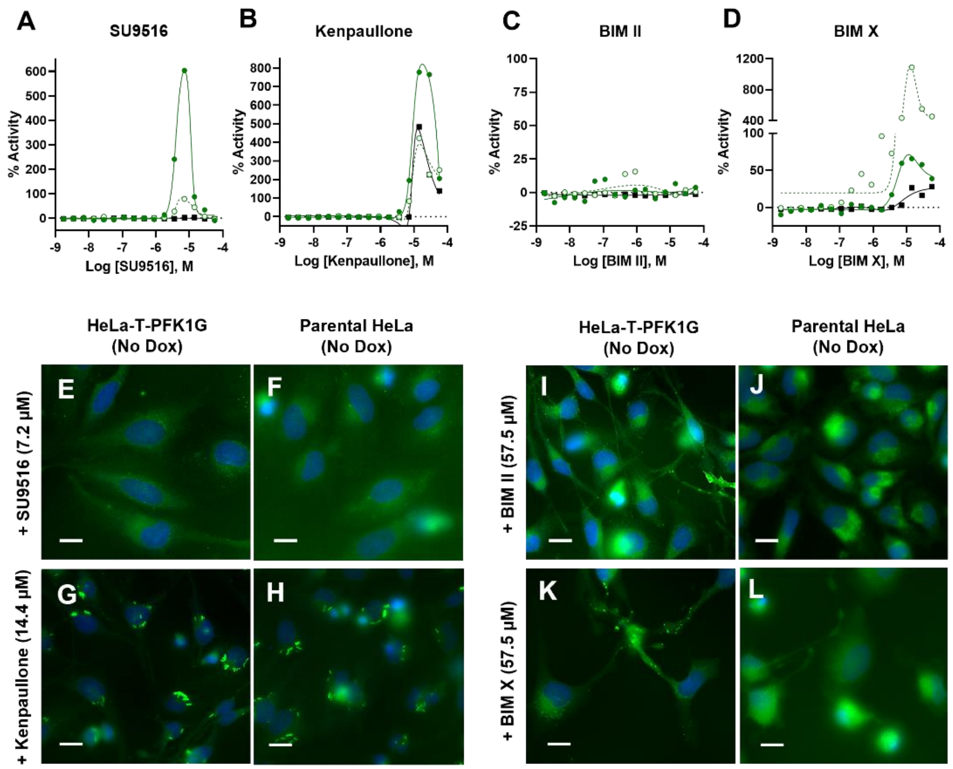


**Supplementary Figure 3. Formation of artifactual fluorescent puncta by chemical compounds.** (**A-D**) Formation of artifactual fluorescent puncta in the presence of SU9516 (**A**), kenpaullone (**B**), BIM II (**C**) or BIM X (**D**) were quantified by an ImageXpress Confocal HT.ai (Molecular Devices) from HeLa-T-PFK1G cells with doxycycline induction (●, solid green circles) or without doxycycline (○, open green circles), and also from parental HeLa cells (■, black squares). Data from HeLa-T-PFK1G cells with or without doxycycline induction were then normalized to DMSO as 0% activity and 11.6 µM SU9516 as 100% activity; while data from parental HeLa cells (■, black squares) were normalized to DMSO as 0% activity and 11.6 µM kenpaullone as 100% activity. GraphPad Prism was used to fit with bell-shaped curves as a function of compound concentration. (**E-L**) Representative images from an ImageXpress imager in the presence of an indicated compound from HeLa-T-PFK1G cells without doxycycline induction (**E**, **G**, **I** and **K**) and from parental HeLa cells (**F**, **H**, **J** and **L**). Note that BIM II and BIM X are control compounds known to cause artifactual fluorescent responses (24). Scale bar, 20 µm.

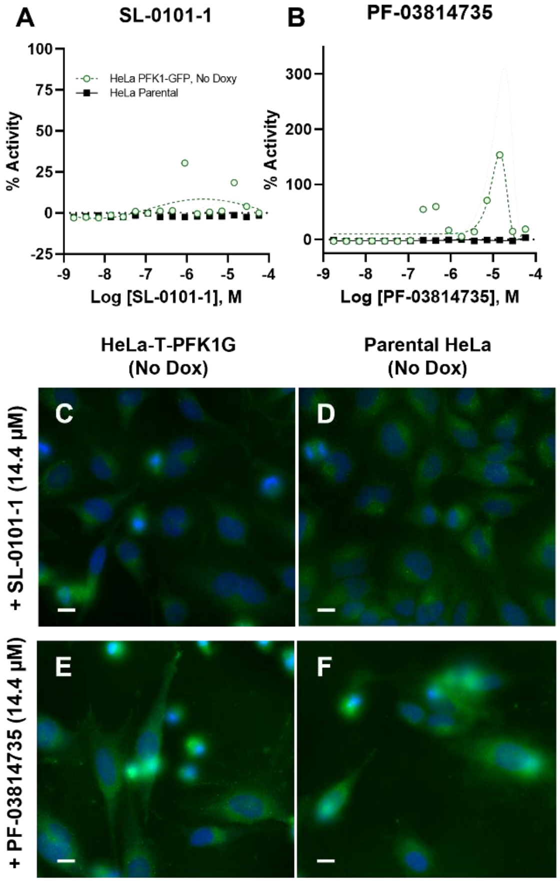


**Supplementary Figure 4. Negligible formation of artifactual fluorescent puncta by the selected kinase inhibitors.** Potential formation of artifactual fluorescent puncta in the presence of SL-0101-1 (**A**) or PF-03814735 (**B**) were investigated by an ImageXpress Confocal HT.ai (Molecular Devices) from HeLa-T-PFK1G cells without doxycycline induction (○, open green circles) as well as from parental HeLa cells (■, black squares). Normalized data to a vehicle control (DMSO) as 0% activity and 11.6 µM SU91516 (Supplementary Figure 3) as 100% activity were plotted in GraphPad Prism and fit with bell-shaped curves as a function of compound concentration. (**C-F**) Representative images from an ImageXpress imager in the presence of an indicated compound from HeLa-T-PFK1G cells without doxycycline induction (**C** and **E**) as well as from parental HeLa cells (**D** and **F**). Scale bar, 20 µm.
